# The song does not remain the same: daily singing of adult songbirds prevents passive changes in song structure independently of auditory feedback

**DOI:** 10.1101/2023.02.22.529516

**Authors:** Daisuke Mizuguchi, Miguel Sánchez-Valpuesta, Yunbok Kim, Ednei B. dos Santos, HiJee Kang, Chihiro Mori, Kazuhiro Wada, Satoshi Kojima

## Abstract

Many songbirds learn to produce songs through vocal practice early in life and continue to sing numerous renditions of their learned songs daily throughout their lifetime. While it is well-known that adult songbirds sing as part of a mating ritual, the functions of singing behavior outside of reproductive contexts are not fully understood. Here, we demonstrate that adult singing outside of reproductive contexts serves to prevent passive changes in song performance. We suppressed the daily singing behavior of adult zebra finches produced in the solo context for two weeks using a reversible behavioral manipulation and examined the effect on song performance. Our results indicated that suppressing daily singing significantly decreased the pitch of song elements and both the amplitude and duration of song motifs. These findings suggest that adult song is not acoustically stable without singing, and that adult birds maintain their song performance by daily singing. Moreover, we found that the changes in song structure caused by singing suppression were substantially recovered within two weeks of free singing, even in deafened birds. Thus, the recovery of song performance does not necessarily require auditory feedback but is predominantly caused by singing behavior per se (i.e., the physical act of singing). Finally, unlike the auditory feedback-dependent song plasticity reported previously, the passive song changes caused by singing suppression were not significantly dependent on age. Taken together, our findings demonstrate that adult songbirds maintain song performance by preventing passive song changes through the physical act of daily singing throughout their life. Such daily singing likely functions as vocal training to maintain the neural and/or muscular system in optimal conditions for song performance in reproductive contexts, similar to how human singers and athletes practice daily to maintain their performance.

## Introduction

Singing behavior is a highly complex motor skill that has been extensively studied across many taxa, including birds, mice, whales, and primates including humans^1^. In songbirds, the primary function of singing is mate attraction and territorial defense in reproductive contexts^2^. Singing behavior in reproductive contexts is observed in the presence of apparent recipients, and thus the purpose of singing can be easily determined by the response of the recipients. In contrast, the functions of singing outside reproductive contexts, produced even when no apparent recipients are around, are much less understood. Although this type of singing behavior has been suggested to play important roles in vocal learning and song structure maintenance^3–5^, the exact function of singing outside reproductive contexts remains controversial^6–9^.

The zebra finch (*Taeniopygia guttata*) is a non-territorial songbird commonly used for neuroethological studies of birdsong. Male zebra finches learn to produce a song during a limited period in early life (closed-ended learner^10^). As adults (>90 days post-hatch [dph]) they repeatedly produce thousands of renditions of the learned song daily throughout life, even when isolated from other birds (i.e., outside reproductive contexts)^11–13^. This daily singing in the solo context, generally called “undirected singing”, has been shown to have a function to refine and maintain song structure: through undirected singing, birds evaluate the song structure using auditory feedback and refine it if necessary^3,4,14–17^ (although other functions of undirected song have been suggested^7–9^). This auditory feedback-dependent function of undirected singing, however, appears to be limited to young adult birds, as the removal of auditory feedback causes substantial changes in song structure in relatively young birds but not in old birds^16,17^. Nevertheless, old birds still sing many renditions of undirected songs every day^12,18^, raising the possibility that undirected singing serves an unknown function in addition to the auditory feedback-dependent refinement of song structure. To explore novel functions of daily undirected singing, we suppressed undirected singing of adult zebra finches for two weeks (Fig. 1*A*) using a method to physically interfere with their singing posture^19^ and examined how their song performance was affected.

**Figure 1.**
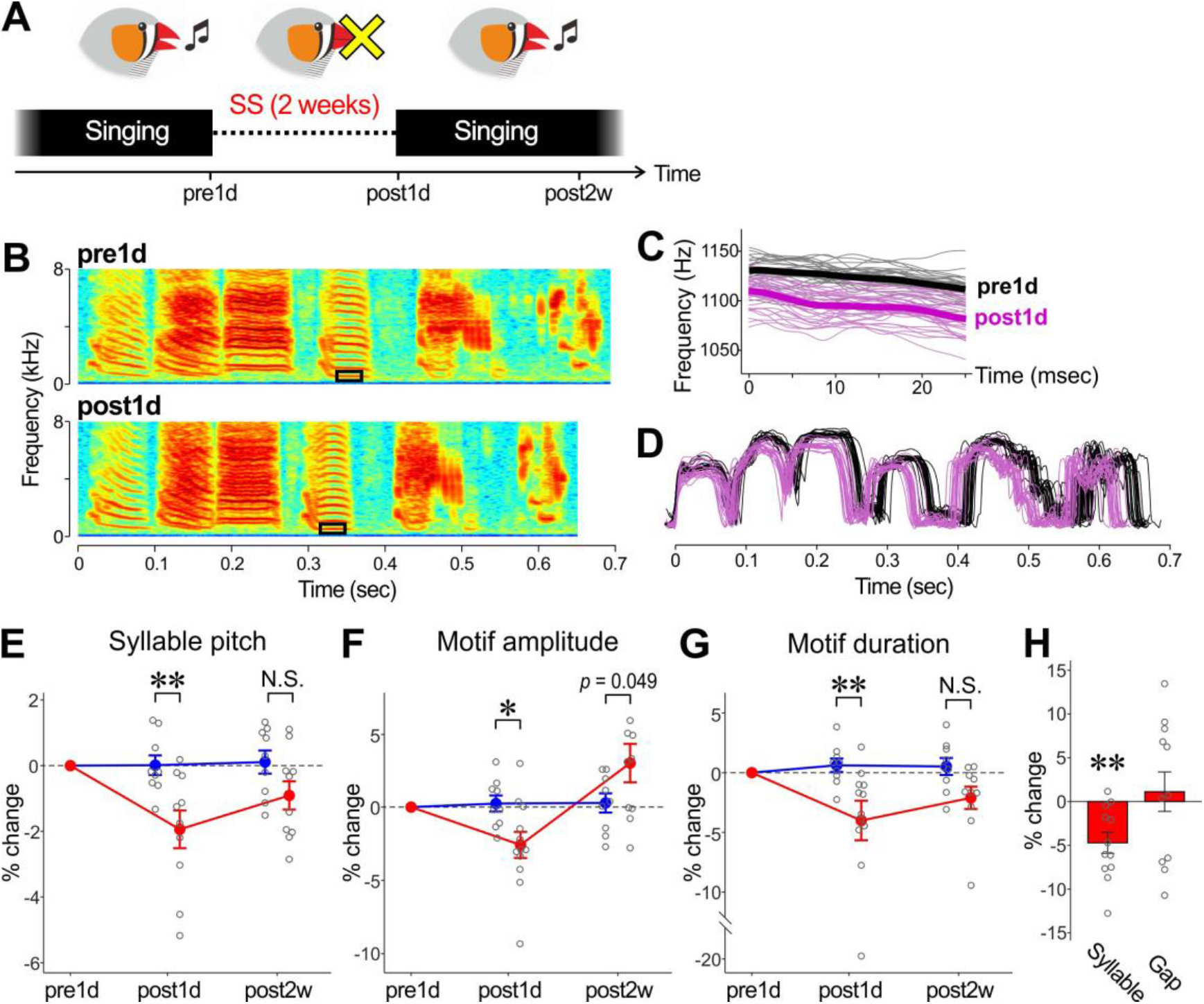
Changes in song performance by suppression of daily undirected singing. (A) Schematic diagram of the experimental procedure. Undirected singing was suppressed for 2 weeks (SS) and songs recorded one day before SS (pre1d) and one day and 2 weeks after SS (post1d and post2w, respectively) were analyzed. (B) Example spectrograms of undirected song motifs at pre1d and at post1d, showing that the overall song structure was maintained. (C) Pitch trajectories of the rectangle regions shown in *B*. Thin lines indicate pitch trajectories from a single syllable rendition; thick lines indicate mean trajectories across all renditions examined; black and magenta lines indicate pre1d and post1d songs, respectively. (D) Overlaid amplitude envelopes of song motifs obtained from the bird shown in *B*. Conversions are as in *C*. (E-G) Percent changes in syllable pitch (E), motif amplitude (F), and motif duration (G) of SS-treated (red lines) and control (blue lines) birds at post1d and post2w relative to pre1d songs. Circles indicate the data of individual birds and lines indicate the mean and SEM across birds. All the song features significantly decreased at post1d in SS birds compared to those in control birds and substantially recovered at post2w (\*\**p* < 0.01, \**p* < 0.05). (H) Percent changes in syllable and silent gap duration of SS-treated birds at post1d relative to pre1d data. Syllable duration but not gap duration significantly changed by SS procedure (\*\**p* < 0.01).

## Results

We compared undirected songs between one day before (pre1d) and one day after (post1d) the singing suppression (SS) period (Fig. 1*A*) in young adult birds (96-129 dph), and found small but significant changes in song structure following the SS period: Although the overall song acoustic structure was maintained (Fig. 1*B*), the pitch of harmonic syllables significantly decreased compared control birds, which were allowed to freely sing for 2 weeks (post1d in Figs. 1*C, E*; n = 10 SS-treated birds and 9 control birds; *p* = 0.002, two-way ANOVA and post-hoc Bonferroni’s multiple comparisons test, see Table 1 for the results of two-way ANOVA). Also, significant decreases were observed in both the root mean square (RMS) amplitude and mean duration of song motifs (a stereotyped sequence of syllables) (post1d in Figs. 1*D, F, G*; *p* = 0.029 for motif amplitude; *p* = 0.003 for motif duration). The decrease in motif duration was predominantly attributable to the decrease in syllable duration rather than in silent gap duration (Fig. 1*H*; *p* = 0.0024 and *W* = 75 for syllable duration and *p* = 0.57 and *W* = 31 for gap duration, Wilcoxon signed-rank test). Moreover, those syllable and motif structures tended to return to their pre-SS levels after 2 weeks of free singing following the SS period, suggesting reversible effects of SS on song structure (post2w in Fig. 1*E-G*; *p* = 0.11 and 0.25 for syllable pitch and motif duration, respectively; the results of motif amplitude were barely significant [*p* = 0.049]). These results provide evidence that neither the syllable nor motif structures of adult songs are stable without daily undirected singing, and suggest that undirected singing has a function to prevent passive changes in song performance to maintain it in the optimal condition.

**Table 1.**
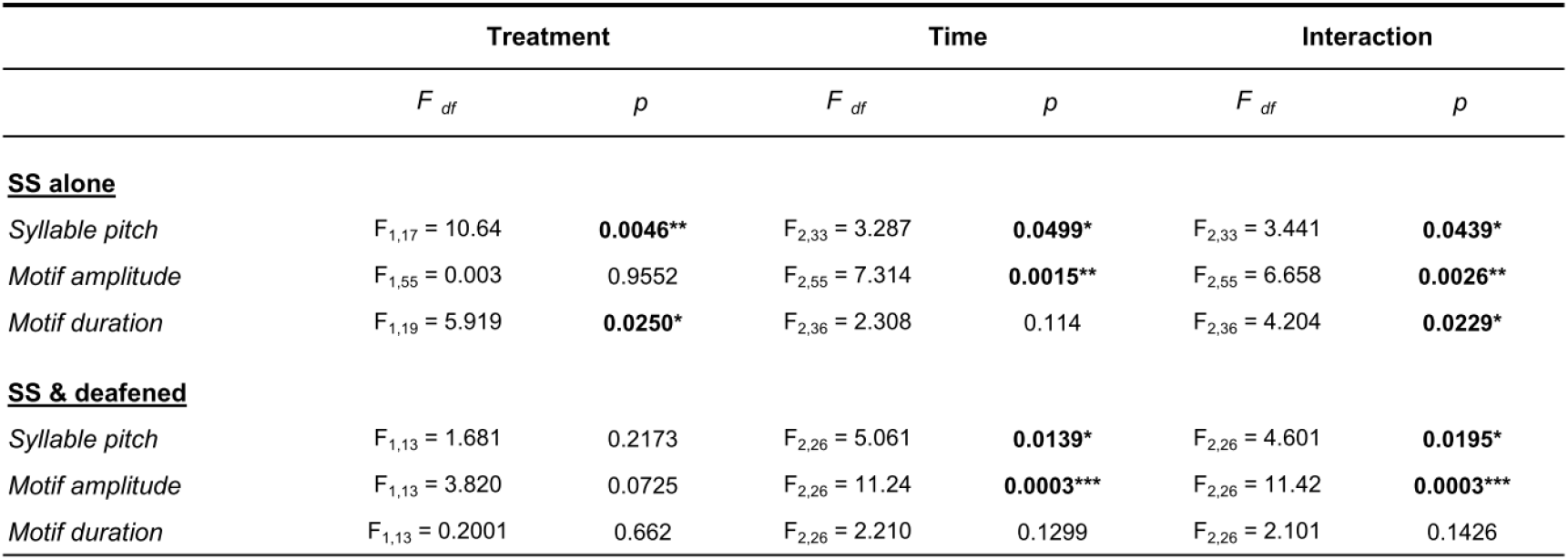
The results of a two-way repeated measures ANOVA analysis for the data in Fig. 1*E-G* (SS alone) and those in Fig. 2*B-D* (SS and deafening).

Undirected singing has been previously shown to function in refinement and maintenance of song structure by providing auditory feedback, through which the birds evaluate their own song structure and correct vocal errors^3,4,14,15^. We wondered whether similar, auditory feedback-dependent mechanisms mediate the recoveries of syllable pitch and motif structure after SS. To examine this, we deafened birds to remove auditory feedback immediately before the SS period and examined whether their song recovery following SS was affected (Fig. 2*A*). Despite the lack of auditory feedback, the deafened birds showed reversible changes in syllable pitch and motif amplitude similar to those observed in intact-hearing birds (Fig. 2*B-C*; n = 6 SS-treated and deafened birds and 9 control [freely singing and intact hearing] birds; for syllable pitch, *p* = 0.013 at post1d and >0.99 at post2w; for motif amplitude, *p* = 0.0001 at post1d and >0.99 at post2w; see Table 1 for the results of two-way ANOVA). A similar trend was observed for motif duration as well, although the results were not significantly different (Fig. 2*D*; *p* = 0.8164 at post1d and 0.2197 at post2w). Also, there was a similar trend for the decreases in syllable duration and silent gap duration as well (Fig. 2*E*), although neither decrease was statistically significant (*p* = 0.44 and *W* = 15 for syllable duration; *p* = 0.69 and *W* = 8 for gap duration). Thus, the recovery of song performance after SS does not necessarily require auditory feedback, and is likely to be caused mainly by singing behavior per se (i.e., the physical act of singing). These results suggest an auditory feedback-independent function of undirected singing to maintain song performance.

**Figure 2.**
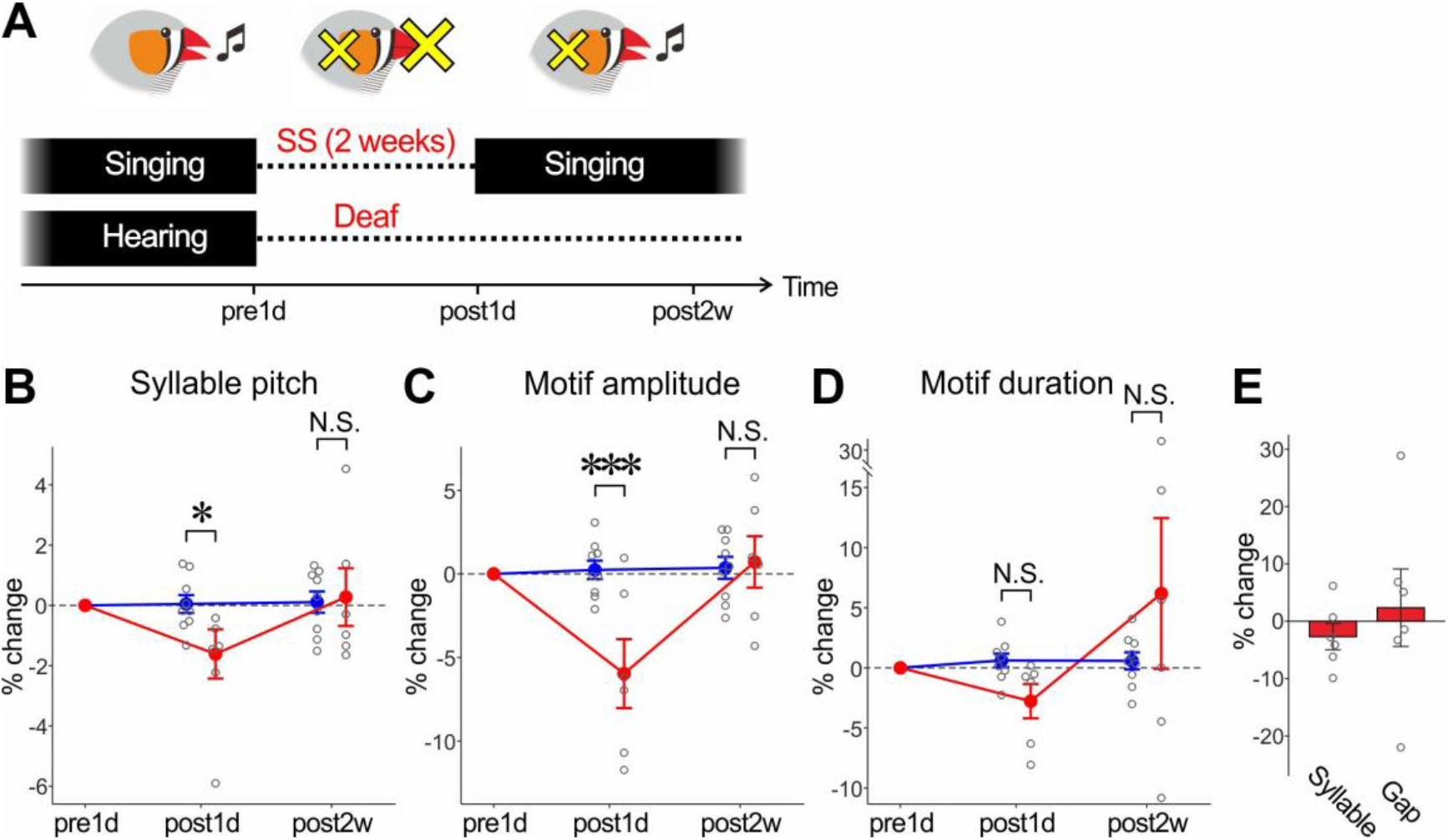
Auditory feedback-independent recovery in song performance after SS through daily undirected singing. (A) Schematic diagram of the experimental procedure. Birds were deafened immediately before SS. (B-D) Percent changes in syllable pitch (B), motif amplitude (C), and motif duration (D) in deafened and SS-treated birds (red lines) and control (normal-hearing and free-singing) birds (blue lines). Conversions are as in Figs. 1*E-G*. \*\*\**p* < 0.001, \**p* < 0.05. (E) Percent changes in syllable and silent gap duration of experimental birds at post1d relative to pre1d data.

In zebra finches, the auditory feedback-dependent function of undirected singing to maintain song structure strongly depends on age: the removal of auditory feedback causes substantial changes in song structure in young adult birds (< ~200 dph) but not in older birds^16,17^. Given that old birds still sing many renditions of undirected songs every day (although the number of undirect songs decreases with age)^12,18^, it is possible that passive changes in song structure can occur independently of age and that those changes are prevented by undirected singing throughout life. To test this possibility, we examined the effect of SS on song structure in relatively old birds with intact hearing (396–809 dph, n = 6 birds) and combined their data with the data of young adult birds shown in Fig. 1*E-G*, to ask whether SS-induced song changes depend on age. In contrast to auditory feedback-dependent song plasticity previously reported^16,17^, we found no significant effects of age on SS-induced song changes (Fig. 3; *r* = −0.018, *p* = 0.94 for syllable pitch; *r* = 0.022, *p* = 0.93 for motif amplitude; *r* = 0.17, *p* = 0.49 for motif duration). Thus, passive changes in adult song structure can occur regardless of age, suggesting that birds produce daily undirected songs to prevent passive song changes throughout life.

**Figure 3.**
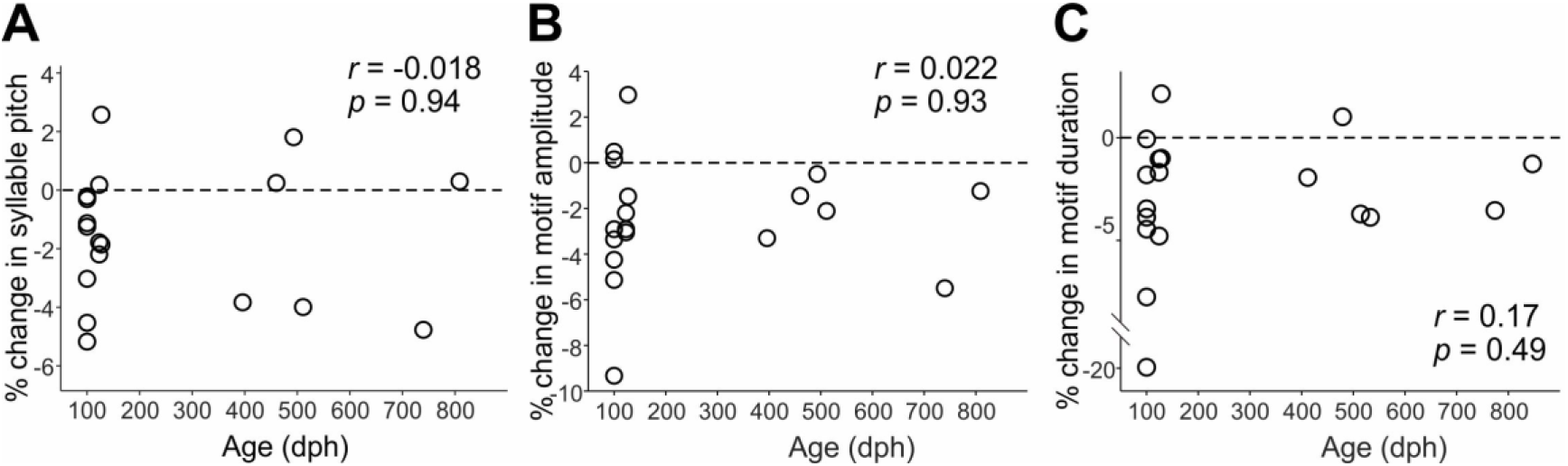
Age-independent changes of the song performance by SS. (A-C) Percent changes in syllable pitch (A), motif amplitude (B), and motif duration (C) at post1d are plotted against the ages of birds. There were no significant effects of age on the changes in the song features.

Because our procedure of SS is likely to induce substantial physical stress in the birds, we cannot exclude the possibility that the passive song changes observed after SS period were predominantly caused by the physical stress induced by the SS procedure rather than the lack of undirected singing. To examine this possibility, we measured plasma corticosterone (CORT) levels in a different group of birds before and during the SS period (see Methods for more detail). We found that many birds increased CORT levels on the first day of the SS period to levels similar to those induced by restraint stress reported in previous studies^20,21^ and subsequently decreased toward the pre-SS levels (Fig. 4*A*; *n* = 10 birds). Also, the increases in CORT levels on the first day of SS period were not significantly correlated with SS-induced changes in the song features that we examined (syllable pitch, motif amplitude, or motif duration) (Fig. 4*B*; *n* = 10 birds). These results support the notion that SS-induced song changes are mainly attributed to the lack of undirected singing rather than the physical stress caused by SS procedure. Moreover, although many birds that received SS exhibited weight loss, an indirect indicator of stress level, no significant correlations were found between the weight changes and the song changes (Fig. 4*C*; the song data in Figs 1 & 2 were replotted to be compared with weight changes), further supporting the independence of SS-induced song changes from physical stress. These results are also consistent with a previous study that the pitch and amplitude of undirected songs are significantly affected by food restriction only when food is restricted in early life but not in adulthood^22^. In summary, the song changes induced by SS are most likely caused by the lack of singing behavior, rather than stress.

**Figure 4.**
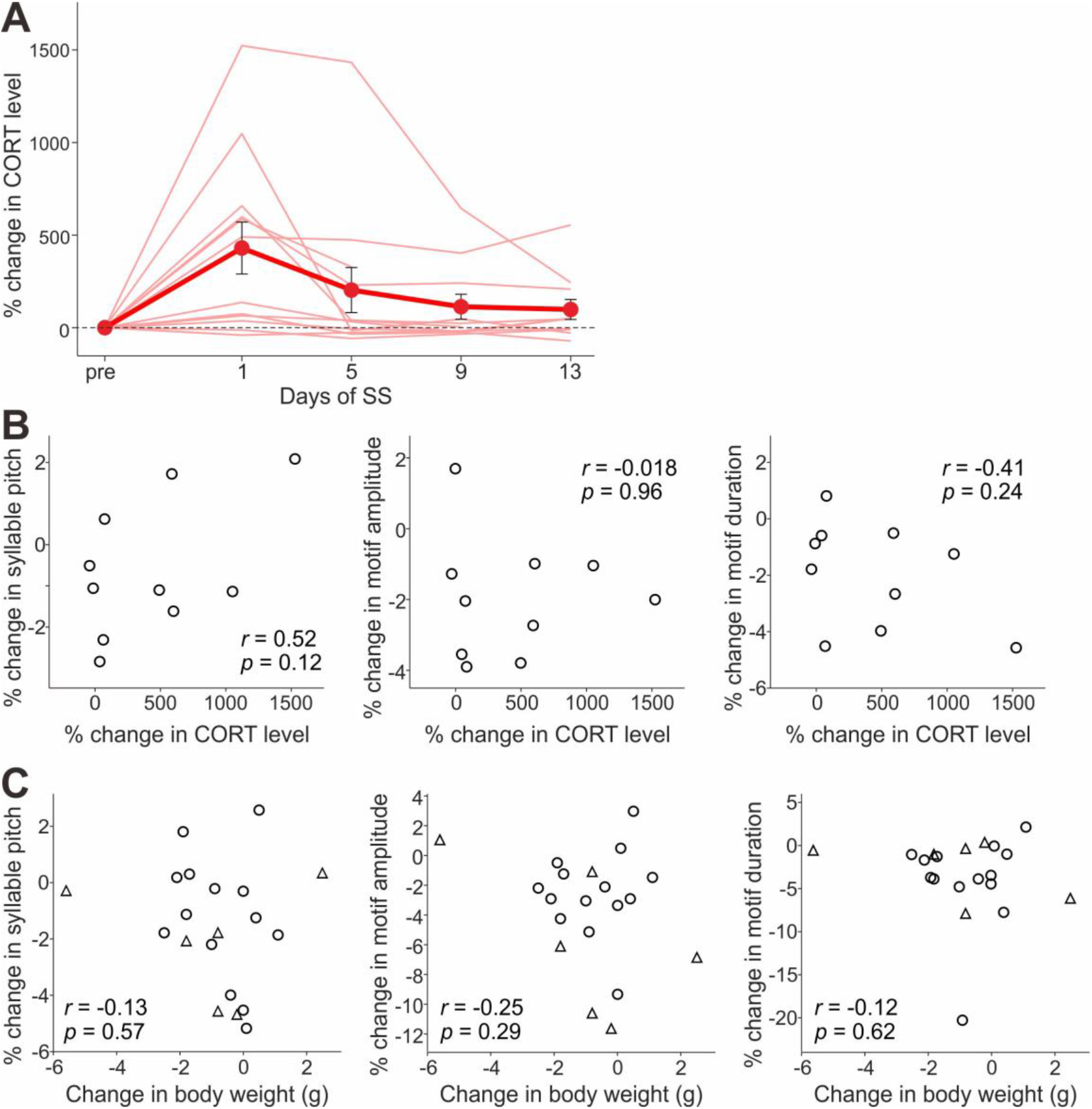
SS-induced changes in song performance are not well explained by stress or bodyweight loss. (A) Percent changes in plasma CORT levels caused by SS treatment; thin lines indicate the data of individual birds and thick lines indicate the mean (±SEM) across the birds. (B) Percent changes in syllable pitch (left), motif amplitude (middle), and motif duration (right), plotted against those in CORT levels measured on the 1^st^ day of SS. (C) Percent changes in syllable pitch (left), motif amplitude (middle), and motif duration (right), plotted against bodyweight changes (the song data in Figs 1 & 2 were replotted). Circles and triangles indicate SS birds with normal hearing and those with hearing impaired (deafened), respectively.

## Discussion

In summary, we found that suppression of undirected singing in adult zebra finches changes both syllable and motif structures of songs, which can be recovered within a short period of free singing. This is the first study in a closed-ended learner showing that the adult song can substantially change over time in the absence of singing behavior, revealing a novel function of undirected singing to prevent passive song changes. Moreover, the recovery of song performance following the SS period does not necessarily require auditory feedback, highlighting the importance of the physical act of singing in the maintenance of adult song performance. Finally, in contrast to song plasticity reported in previous studies^16,17^, SS-induced song changes do not depend on age, providing a plausible explanation for why adult birds sing many renditions of the undirected song throughout life even in old age. Taken together, our findings provide evidence that zebra finches maintain their song performance by preventing passive song changes through the physical act of daily singing throughout life. Because male zebra finches produce courtship songs with the structure that is virtually the same as that of undirected song^10^, it is likely that they maintain the song production system in the optimal condition through undirected singing to prepare for the best song performance in reproductive contexts.

We previously showed that temporary suppression of undirected singing (for 5 or 10 h) in adult zebra finches induces intense singing immediately after the SS period, indicating that intrinsic motivation for undirected singing increases while singing is suppressed^23^. This SS-induced increase in singing motivation and resulting intense singing can be explained by the song maintenance function of undirected singing that we found in the present study. It is reasonable to assume that birds produce intense singing after SS to compensate for the loss of vocal practice during SS and to quickly re-optimize the vocal system and song structure for future courtship activity. Although it remains unclear whether adult song structure actually deteriorates during such short SS periods and recovers through subsequent intense singing, substantial song deterioration during non-singing periods and improvement during subsequent singing are observed in juvenile birds producing immature songs^11,13^. Moreover, this idea of vocal re-optimization through intense singing after SS is consistent with a hypothesized role of the “dawn chorus”, intense singing behavior at dawn widely observed in many songbird species^24,25^. Because dawn chorus occurs after a long break of daytime singing by the night and is accompanied by significant improvement in song structure^26,27^, it has been hypothesized that through intense dawn singing, birds warm up their vocal system to prepare for future courtship and territorial defense activities just as human athletes and singers do before their performance^26,27^. Our results in the present and previous studies provide empirical support for this warm-up hypothesis of dawn chorus, advancing our understanding of the functional roles of non-reproductive singing in wild birds as well as in captive birds.

While our findings suggest that the limited singing behavior of adult birds results in substantial changes in the song performance, the physiological mechanisms underlying such passive song changes should be further studied. It is presumably caused, at least in part, by weakening (atrophy) of the syringeal muscles responsible for song production, similar to the atrophy of skeletal muscles that is generally known to be caused by a lack of physical activity. Comparing the syringeal muscle volumes or fiber sizes before and after SS would be necessary to examine any possible muscle atrophy. Given the decreasing effect of SS on motif and syllable duration, it is also possible that the neural circuitry that generates song structure passively drifts without daily undirected singing. Consistent with this idea, substantial changes in singing-related neural activity in the song motor pathway occur primarily during the nighttime (when birds do not sing), and less frequently during the daytime^28,29^. Also, changes in motif duration are observed even in birds that freely sing when the central neural circuits are manipulated^30,31^, further supporting the possibility that SS-induced song changes are attributable, at least in part, to passive changes in neural circuits. Comparing singing-related neural activity patterns before and after SS would be necessary to examine this possibility. Regardless of the mechanisms of passive song changes induced by the lack of daily undirected singing, our findings open the possibility that the physical activity-dependent and hearing-independent mechanisms for maintaining vocal performance are commonly used in many other bird species and other animal taxa, including non-vocal learning birds and mammals.

## Materials and Methods

### Subjects

Adult male zebra finches (98–809 dph) were bred in our colony at Korea Brain Research Institute and individually housed in sound-attenuating chambers (MC-050, Muromachi Kikai, or custom-made ones) on a 14:10-h light:dark cycle throughout the experiments. Their care and treatment were reviewed and approved by the Animal Care and Use Committee at Korea Brain Research Institute.

### Song recording

Songs were recorded using a microphone (PRO35, Audio-Technica) positioned 5~10 cm above the cage (20 x 20 x 20 cm) and a custom-made recording program at a 32-kHz sampling rate with 16-bit resolution. To minimize the influence of the bird’s position on the amplitude of song recording, no perch was placed in the cage and the food was provided in a petri dish placed on the bottom of the cage so as to have the bird sing at the cage bottom (zebra finches rarely sing while holding on to the side walls or ceiling of the cage). Song recording was triggered when the program detected 4 or 5 successive song elements, each of which reached predefined thresholds of amplitude and duration, and ended when a silent period lasted >0.5 sec. Our recording system thus captured mostly the song bouts, although short calls (and cage noises) were occasionally recorded. We counted the number of song bouts daily to monitor singing activity (Supplementary Tables 1-2). Birds that produced songs containing at least one harmonic syllable with sufficient singing rates (>200 undirected song bouts per day) were used for experimentation.

### Singing suppression

Spontaneous undirected singing was suppressed for two weeks using a previously published method^19^. Briefly, a custom-made weight was attached to the neck of the bird each morning to interfere with his singing posture to prevent him from singing (the weight shifted the posture toward an inferior position). The weight (20.1-32.6 g) was adjusted daily for each bird so that the procedure mostly suppressed undirected singing but did not severely affect the daily behaviors of the bird, such as eating, drinking, preening, and calling (body weight and health conditions were carefully monitored every day). Each night, the weight was removed when the light was turned off as no singing behavior was observed during the night. With this method, the daily singing amount (the number of song bouts per day) decreased to less than 10% of that of pre-SS (see Supplementary Tables 1-2). Birds in the control group were allowed to sing freely without a weight all time.

### Song analysis

To examine the effects of SS on song performance, we compared the acoustic and temporal structure of songs recorded 1 day before SS (pre1d) with songs recorded 1 day and 2 weeks after SS (post1d and post2w, respectively; Fig. 1 *A*). Adult zebra finch’s songs consist of stereotyped sequences of several syllables, which are termed “motifs,” in which syllables are separated by silent gaps (see Fig. 1*B*). To test the effect of SS on the pitch of individual syllables, >30 motifs from pre1d, post1d, and post2w songs were randomly selected for each bird, and syllables containing clear harmonic structure were extracted. For each syllable rendition, the pitch trajectory was obtained from a segment of a sound waveform described in a previous study^31,32^. Briefly, spectrograms were calculated using a Gaussian-windowed short-time Fourier transform (*σ* = 1 ms) sampled at 8 kHz, and a pitch trajectory was obtained by calculating the fundamental frequency in individual time bins. The fundamental frequency was calculated by parabolic interpolation. For each syllable, all renditions of pitch trajectories were aligned by the syllable onsets, which were determined based on amplitude threshold crossings, and the mean pitch values over the flat portions of pitch trajectories were averaged across all renditions. All the birds examined in the present study had at least one syllable with a clear harmonic structure in their song motif; for the birds that had more than one harmonic syllable, we calculated the effects (percent changes) of SS on the pitch of individual syllables and averaged them across the syllables for each bird (the mean number of syllables examined per bird = 1.76; max = 3).

For the amplitude measurements, 12-20 motifs with low background noise levels from pre1d, post1d, and post2w songs were randomly selected for each bird. We then measured the RMS amplitude values for individual motifs and averaged them. Since we were only interested in SS-induced changes in the motif amplitude, we did not calculate the absolute amplitude and instead obtained percent changes relative to the amplitude of pre1d song. Because our cage setting limited birds’ singing sites to the bottom of the cage (see the description of song recording above), the distance between a singing bird and the microphone did not dramatically changes, and therefore changes in the RMS amplitude across different time points are attributed mainly to changes in song volume rather than changes in singing sites. One control bird was excluded from the amplitude analysis since the amplitude gain was not consistent over the song recording due to the malfunction of the recording system.

Durations of song motifs, syllables, and silent gaps were measured by thresholding amplitude envelopes that were normalized to their peak amplitudes (the same thresholds were used for all the time points for each bird). The obtained song segmentations were visually inspected and manually modified if necessary, so that syllable/gap segmentation patterns were identical across motif renditions. Percent changes in syllable duration or in silent gap duration were calculated for individual syllables or gaps, and then averaged across different syllables or gaps for each bird.

### Deafening

For experiments to test whether auditory feedback was related to the song change after the SS period, we deafened young adult birds immediately before SS by bilateral cochlea removal following a previously published method^5,18^ and examined whether song recovery following SS was affected. SS-induced changes in syllable pitch, motif amplitude and motif duration were analyzed in the same way as normal-hearing birds. One bird was excluded from the analysis because the song structure was severely deteriorated by deafening.

### Corticosterone analysis

To measure the stress levels of SS-treated birds, blood samples were collected (<75 μl for each sampling) at 5-time points before and during the SS period (pre4d, post1d, post5d, post9d, and post13d) by puncturing the brachial wing vein with a sterile needle (23G) 1 hour after the weight attachment. The blood samples were then centrifuged at 3500 rpm for 20 min at room temperature (approximately 20 °C). The separated serums were stored at −20 °C until the assay. Corticosterone (CORT) levels were measured using an Enzyme Immunoassay kits (Enzo Life Science). Two separate 96-well plates were used in this study, and a separate standard curve was run on each plate. All serum samples (1:20 dilution) and standards were run in duplicates. Plates were read on Absorbance 96 plate reader (Enzo Life Sciences). CORT levels were determined from 4-PL regression of 5 standards ranging from 32 to 20,000 pg/ml. We calculated intra-assay variability as the average %CV between duplicates for each assay and inter-assay variability as %CV among CORT concentrations calculated from each standard curve for 5 corticosterone standards run in each assay. Average intra- and inter-assay variability was 18.0% and 4.8%, respectively.

### Statistics

Statistical tests were performed using R 3.6.2 (R Core Team and R Foundation for Statistical Computing) or MATLAB (MathWorks). The effects of SS on syllable pitch and motif amplitude and duration were examined using a two-way ANOVA and post-hoc Bonferroni’s multiple comparisons test. The effects of SS on syllable duration and silent gap duration were examined using a Wilcoxon signed-rank test. The relationships of song changes with age, CORT changes, and weight changes were examined using Pearson’s correlation coefficients.

## Data availability

All data are included in the article and supporting information.

## Acknowledgments and funding sources

This work was supported by grants from Korea Brain Research Institute basic research program through Korea Brain Research Institute funded by Ministry of Science and ICT (Grant 23-BR-01-01 to S.K.). We thank Dr. Ji Young Mun, and Dr. Min Kyo Jung at Korea Brain Research Institute and Dr. Ryosuke O. Tachibana at Tokyo University for their valuable comments and discussion.

