## Supplementary Tables 1-2 for "The song does not remain the same: daily singing of adult songbirds prevents passive changes in song structure independently of auditory feedback"

### Supplementary Materials

**Supplementary Table 1.** Daily counts of the number of undirected song bouts before and during SS in intact hearing birds. Data of the birds used for Figs. 1 & 3 are shown.

| Manipulation | SS (intact hearing) | | | | | | | | | | | | | | | |
|---|---|---|---|---|---|---|---|---|---|---|---|---|---|---|---|---|
| **Bird ID** | b17p6 | b19r17 | m67y50 | y69r26 | y92w65 | w78w6 | m52m41 | b80p39 | m56p64 | w32r84 | o26b44 | w51o37 | r47w46 | b81k6 | m5k36 | y1r60 |
| **Age (dph)** | 100 | 100 | 100 | 100 | 100 | 100 | 100 | 123 | 123 | 123 | 396 | 460 | 493 | 511 | 740 | 809 |
| ***Before SS*** | | | | | | | | | | | | | | | | |
| **pre1d** | 1602 | 1403 | 558 | 821 | 1753 | 1938 | 1850 | 1089 | 1427 | 1143 | 205 | 734 | 633 | 362 | 536 | 304 |
| ***During SS*** | | | | | | | | | | | | | | | | |
| **day1** | 0 | 0 | 69 | 0 | 0 | 0 | 0 | 0 | 0 | 0 | 0 | 0 | 0 | 0 | 41 | 0 |
| **day2** | 31 | 165 | 2 | 447 | 0 | 0 | 0 | 1 | 0 | 4 | 0 | 1 | 3 | 0 | 4 | 0 |
| **day3** | 0 | 284 | 0 | 556 | 0 | 0 | 0 | 121 | 22 | 300 | 19 | 0 | 11 | 0 | 0 | 0 |
| **day4** | 0 | 0 | 0 | 44 | 4 | 0 | 0 | 8 | 4 | 0 | 0 | 0 | 62 | 9 | 0 | 0 |
| **day5** | 0 | 0 | 0 | 0 | 0 | 0 | 0 | 0 | 2 | 3 | 0 | 0 | 0 | 2 | 0 | 1 |
| **day6** | 0 | 116 | 0 | 0 | 1 | 0 | 0 | 246 | 24 | 1 | 0 | 0 | 0 | 0 | 0 | 1 |
| **day7** | 0 | 0 | 0 | 16 | 1 | 0 | 0 | 0 | 2 | 1 | 0 | 0 | 2 | 0 | 0 | 0 |
| **day8** | 0 | 0 | 0 | 17 | 0 | 55 | 0 | 0 | 53 | 0 | 0 | 0 | 0 | 2 | 0 | 0 |
| **day9** | 0 | 0 | 0 | 0 | 0 | 15 | 0 | 500 | 0 | 5 | 0 | 2 | 1 | 0 | 0 | 0 |
| **day10** | 0 | 0 | 0 | 0 | 0 | 15 | 0 | 0 | 3 | 0 | 0 | 0 | 0 | 0 | 0 | 1 |
| **day11** | 0 | 0 | 6 | 1 | 0 | 31 | 0 | 0 | 241 | 0 | 0 | 0 | 0 | 0 | 0 | 0 |
| **day12** | 0 | 0 | 0 | 0 | 0 | 4 | 0 | 0 | 0 | 0 | 0 | 0 | 1 | 0 | 0 | 0 |
| **day13** | 0 | 0 | 1 | 0 | 0 | 2 | 0 | 0 | 0 | 0 | 0 | 1 | 0 | 0 | 0 | 0 |
| **day14** | 0 | 0 | 0 | 0 | 0 | 2 | 0 | 0 | 238 | 0 | 0 | 0 | 0 | 0 | 0 | 0 |
| **Mean during SS (/day)** | 2.2 | 40.4 | 5.6 | 77.2 | 0.4 | 8.9 | 0.0 | 62.6 | 42.1 | 22.4 | 1.4 | 0.3 | 5.7 | 0.9 | 3.2 | 0.2 |
| **% decrease from pre1d** | 99.9 | 97.1 | 99.0 | 90.6 | 100.0 | 99.5 | 100.0 | 94.3 | 97.1 | 98.0 | 99.3 | 100.0 | 99.1 | 99.7 | 99.4 | 99.9 |

**Supplementary Table 2.** Daily counts of the number of undirected song bouts before and during SS in deafened birds. Data of the birds used for Fig. 2 are shown.

| **Manipulation** | **SS & deafened** | | | | | |
|---|---|---|---|---|---|---|
| **Bird ID** | **w27k27** | **o87r58** | **k6y57** | **k83w25** | **p1r56** | **y34m86** |
| **Age (dph)** | 98 | 99 | 100 | 100 | 100 | 101 |
| ***Before SS*** | | | | | | |
| **pre1d** | 265 | 1167 | 1089 | 1489 | 2825 | 1965 |
| ***During SS*** | | | | | | |
| **day1** | 0 | 0 | 0 | 0 | 0 | 0 |
| **day2** | 0 | 0 | 0 | 0 | 0 | 0 |
| **day3** | 0 | 0 | 0 | 0 | 0 | 0 |
| **day4** | 0 | 0 | 0 | 0 | 16 | 0 |
| **day5** | 31 | 0 | 0 | 0 | 0 | 43 |
| **day6** | 3 | 0 | 0 | 26 | 15 | 0 |
| **day7** | 101 | 3 | 0 | 0 | 0 | 86 |
| **day8** | 33 | 141 | 1 | 1 | 0 | 27 |
| **day9** | 0 | 0 | 1 | 0 | 0 | 0 |
| **day10** | 0 | 0 | 2 | 0 | 0 | 0 |
| **day11** | 0 | 0 | 1 | 0 | 0 | 0 |
| **day12** | 0 | 0 | 3 | 0 | 0 | 0 |
| **day13** | 9 | 0 | 0 | 0 | 0 | 0 |
| **day14** | 2 | 0 | 0 | 0 | 0 | 0 |
| **Mean during SS (/day)** | 12.8 | 10.3 | 0.6 | 1.9 | 2.2 | 11.1 |
| **% decrease from pre1d** | 95.2 | 99.1 | 99.9 | 99.9 | 99.9 | 99.4 |
